# HPC-Atlas: Computationally Constructing A Comprehensive Atlas of Human Protein Complexes

**DOI:** 10.1101/2023.01.03.522554

**Authors:** Yuliang Pan, Ruiyi Li, Wengen Li, Liuzhenghao Lv, Jihong Guan, Shuigeng Zhou

## Abstract

A fundamental principle of biology is that proteins tend to form complexes to play significant roles in the core functions of cells. For a complete understanding of human cellular functions, it is crucial to have a comprehensive atlas of human protein complexes. Unfortunately, we still lack such a comprehensive atlas of experimentally validated protein complexes, which prevents us from gaining a complete understanding of the compositions and functions of human protein complexes and biological mechanisms. To fill this gap, we built HPC-Atlas, as far as we know, the most accurate and comprehensive atlas of human protein complexes available to date. We integrated two latest protein interaction networks, and developed a novel computational method to identify nearly 9000 protein complexes, including many previously uncharacterized complexes. Compared with the existing works, our method achieves outstanding performance on both test and independent sets. Furthermore, with HPC-Atlas we identified 751 SARS-CoV-2 affected human protein complexes, and 456 multifunctional proteins that contain many potential moonlighting proteins. These results suggest that HPC-Atlas can serve as not only a computing framework to effectively identify biologically meaningful protein complexes by integrating multiple protein data sources, but also a valuable resource for exploring new biological findings. The HPC-Atlas webserver is freely available at http://www.yulpan.top/HPC-Atlas.

## Introduction

Protein complexes are composed of proteins that interact with each other, they carry out many essential functions of cells, including replication, transcription, and protein degradation [1,2]. Comprehensive characterization of their compositions can provide critical insights into cellular functions and facilitate the understanding of disease-related pathways [3,4]. Related studies have shown that the human genome contains more than 20,000 genes, which encode tens of thousands of different proteins in human cells [5]. It has been estimated that over 80% human proteins participate in complexes [5]. The comprehensive resource of mammalian protein complexes (CORUM) [6] has been widely used as the gold standard for human protein complexes. However, the latest version of CORUM (version 3.0) contains only 2916 human protein complexes covering 3674 different proteins. Hence, we still have very limited knowledge of protein complexes in human cells.

Although many computational approaches have been developed to identify protein complexes [7−16], most of them were designed mainly based on the saccharomyces cerevisiae protein interaction network (PIN). Compared with the human protein interaction network, the saccharomyces cerevisiae network is smaller, and thus it is relatively easy to identify the complexes from this network. Moreover, these approaches generally consider the densely connected subgraphs in the PIN as complexes. However, the human PIN is larger and sparser, and many human protein complexes are small complexes composed of two or three proteins, which poses serious challenges to the traditional identification approaches. Recently, Drew et al. [17] developed a machine learning pipeline to identify 6965 human protein complexes from integrated protein interaction data of over 15,000 proteomic experiments. But this integrated experimental dataset still contains only limited human proteins, and they did not develop any new complex identification algorithm (they used existing methods) for this dataset or network.

In the past two decades, high-throughput techniques, such as yeast 2-hybrid (Y2H) [18] and affinity purification with mass spectrometry (AP-MS) [19], have significantly increased the coverage of protein interactions across the human proteome [20]. Two recently released reference maps of human protein interactome (HuRI [21] and BioPlex [22]) generated by Y2H and AP-MS respectively have substantially advanced this field. Although these techniques have identified a large number of protein interactions, the coverage of the entire human interactome is still limited. Despite the fact that these techniques tend to explore different parts of the human proteome, the sets of identified interactions only partially overlap. This provides an opportunity to integrate the two reference maps and subsequently build a more comprehensive human protein interactome, from which more protein complexes can be identified via computational methods.

In this paper, we present HPC-Atlas (**H**uman **P**rotein **C**omplexes **Atlas**), to the best of our knowledge, the most accurate and comprehensive atlas of human protein complexes to date, which contains nearly 9000 identified protein complexes across 16,632 proteins, including many previously uncharacterized protein complexes. HPC-Atlas was built by a new and effective computational framework over a more comprehensive protein interaction network by integrating two different latest protein interaction networks. Our experimental results show that the proposed method outperforms 15 state-of-the-art protein complex identification methods. Further research revealed that the previously uncharacterized protein complexes in HPC-Atlas have high complex scores, and are strongly supported by significantly enriched functional annotations. Among them, there are 751 complexes related to SARS-CoV-2, and 456 multifunctional proteins that may be crucial moonlighting proteins, participating in multiple complexes. These results show that our atlas covers substantially more human proteins and contains much more biologically meaningful complexes.

## Results

### HPC-Atlas overview

This work constructed a comprehensive atlas of human protein complexes by first integrating two latest protein interaction datasets or networks (HuRI [21] and BioPlex [22]) to get a more comprehensive protein interaction network, and then developing a customized complex identification method, which was finally applied to the integrated PIN, as shown in **Figure 1**. First, we expanded and integrated the two existing protein interaction networks (PINs) of the latest version to construct a larger one (Figure 1A). As each existing PIN contains different protein interactions, the combination of them can create a more comprehensive PIN. Then, we calculated a feature vector for each pair of proteins by merging several different types of features, including subcellular localization, position-specific scoring matrix (PSSM) based features, gene ontology (GO) semantic similarity, protein chain length, and protein domain interactions (Figure 1B). The resulting PIN is called *PIN of featured edges* (FE-PIN in short). Furthermore, we classified edges in the network into two types: *edges within complexes* (denoted by c-edges, meaning complex edges) and *edges outside complexes* (denoted by nc-edges, meaning non-complex edges). Concretely, c-edges are those whose two end proteins simultaneously belong to at least one complex, and nc-edges are those whose two end proteins do not lie in any complex simultaneously. We trained a deep forest classifier [23] to label the edges in FE-PIN by using training data from the gold standard CORUM set. With this classifier, all edges without labels previously can be labeled: each edge without label is assigned a label probability, indicating how possibly the edge is a c-edge or an nc-edge. Thus, we got a *PIN of labeled edges*, which is denoted by LE-PIN in short (Figure 1C). Protein complexes were subsequently identified from LE-PIN by a specifically designed protein complex detection algorithm based on the c-edges. The final set of protein complexes was generated by ranking the identified complexes via their scores, which was further used for exploring new complexes, multifunctional proteins, and COVID-19 related complexes (Figure 1D).

**Figure 1.**
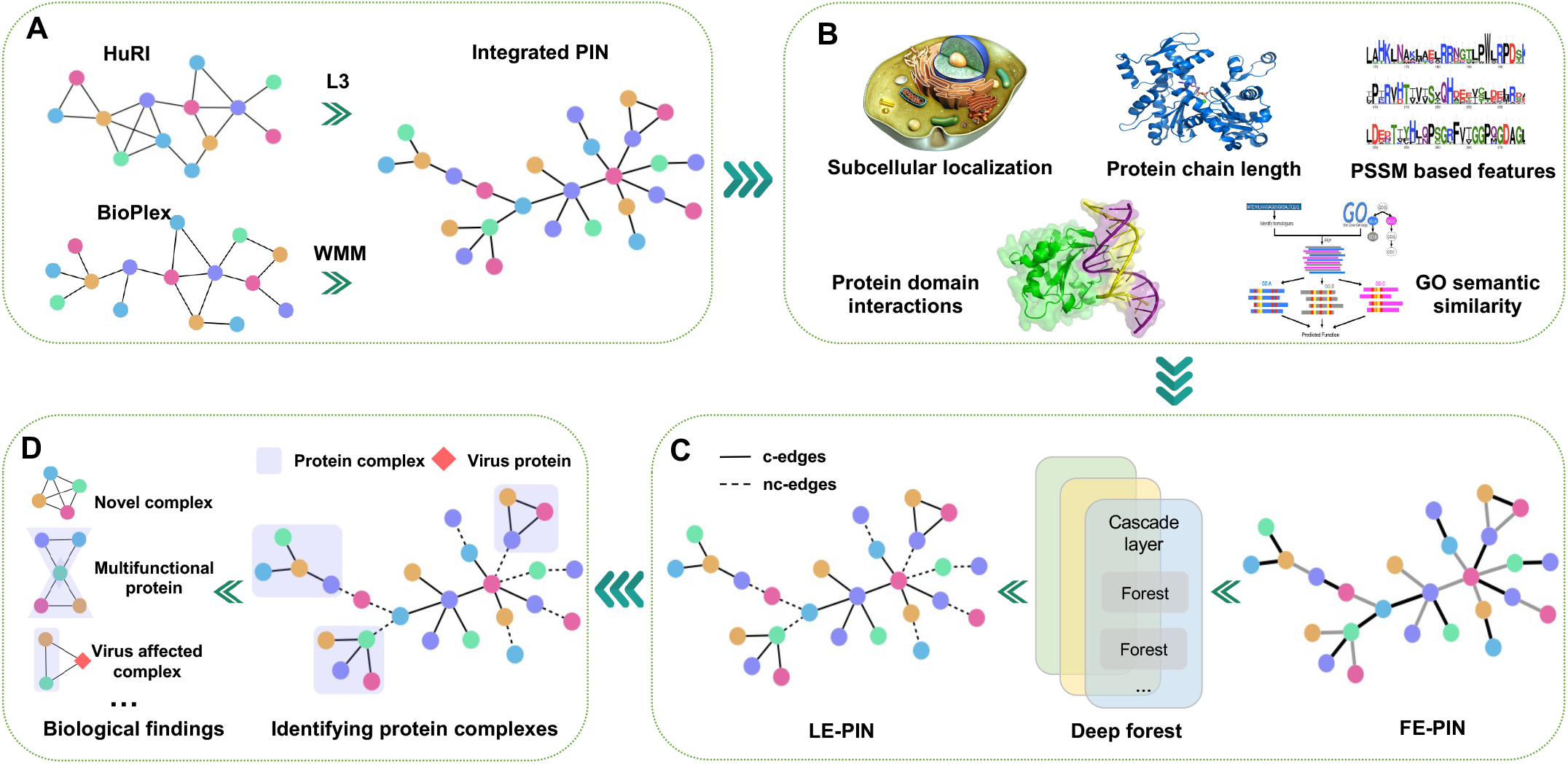
The pipeline of HPC-Atlas. **A**. Two reference PINs HuRI and BioPlex were combined into an integrated PIN after being expanded with predicted interactions by the L3 [24] and WMM [25] algorithms, respectively. **B**. Calculating the features of all protein-protein interactions (or PIN edges), including subcellular localization, PSSM based features, GO semantic similarity, protein chain length and protein domain interactions, which leads to the FE-PIN, i.e., PIN of featured edges. In FE-PIN, different colors of edges correspond to different features. **C**. Training a deep forest classifier to label the edges in FE-PIN, which results in the LE-PIN, i.e., PIN of labeled edges. LE-PIN is actually a weighted PIN. Here, solid edges are c-edges with larger weights, and dashed edges are nc-edges with smaller weights. **D**. A new method was developed to identify complexes from the LE-PIN, and thus a comprehensive atlas of human protein complexes was constructed, which was further used for biological discoveries. PIN, protein interaction network; L3, Length three; WMM, weighted matrix model; PSSM, Position-specific scoring matrix; GO, Gene ontology; c-edges, complex edges; nc-edges, non-complex edges.

### Integrating two human PINs (HuRI and BioPlex)

In general, there is very low overlap between current PINs due to employing different high-throughput techniques [17,26]. Therefore, we combined two recently released PINs HuRI and BioPlex to construct a larger protein interaction map. The HuRI network was generated by Y2H experiments, which consists of 63,132 protein-protein interactions (PPIs) involving 8985 proteins. The BioPlex network was derived from AP-MS experiments of 239T and HCT116 cells, comprising 167,932 PPIs across 14,484 proteins.

By comparing the two PINs, we found that the overlapping ratios of proteins between them are 47% (BioPlex overlapping HuRI) and 76% (HuRI overlapping BioPlex) respectively. However, the overlap ratios of PPIs are much lower, only 1% (BioPlex overlapping HuRI) and 2.6% (HuRI overlapping BioPlex). Here, given two PINs A and B, the ratio of A overlapping B is evaluated as |*A* ∩ *B*| / |*A*|, |•| means the number of nodes or edges in a PIN. There are four possible reasons of the low overlapping ratios between different large-scale PINs. Firstly, the principles of PPI testing by Y2H and AP-MS experiments are similar to the spoke model (**Figure 2A**). Since only the interactions between bait proteins and their prey are considered, all true prey-prey interactions are ignored. Secondly, different research teams constructed large-scale PINs using different cell types and proteins. Thirdly, the existing experimental methods tend to find specific types of PPIs, for example, membrane and soluble protein interactions. Last but not least, experimental methods inevitably generate false positive interactions. To recover false negative PPIs and increase the coverage of the interactome, we applied length three [24] (L3) and weighted matrix model [25] (WMM) algorithms to identify new high confidence PPIs from HuRI and BioPlex respectively. Previous works [17,24,26] have demonstrated that these two methods can effectively find false negative PPIs of high confidence. Then, we integrated the two PINs after adding the predicted PPIs to build a more comprehensive human PIN, which contains 2,658,160 PPIs over 16,632 proteins. For comparison, we present the statistics of the two original PINs and the integrated PIN in Table S1. Compared with HuRI and BioPlex, the integrated PIN captures much more PPIs, which greatly improves the coverage of the human protein interactome. For example, KDM1A interacts with HDAC2, and H3-3B interacts with HAT1, which have been verified in the STRING [27] database, and successfully identified and added to our integrated PIN. We further analyzed the degree distribution of protein nodes in different PINs (see Figure S1) and found that the degree distribution of protein nodes is similar to the scale-free network, that is, a few proteins interact with a large number of proteins, while most proteins in the PIN interact with only a small number of proteins.

**Figure 2.**
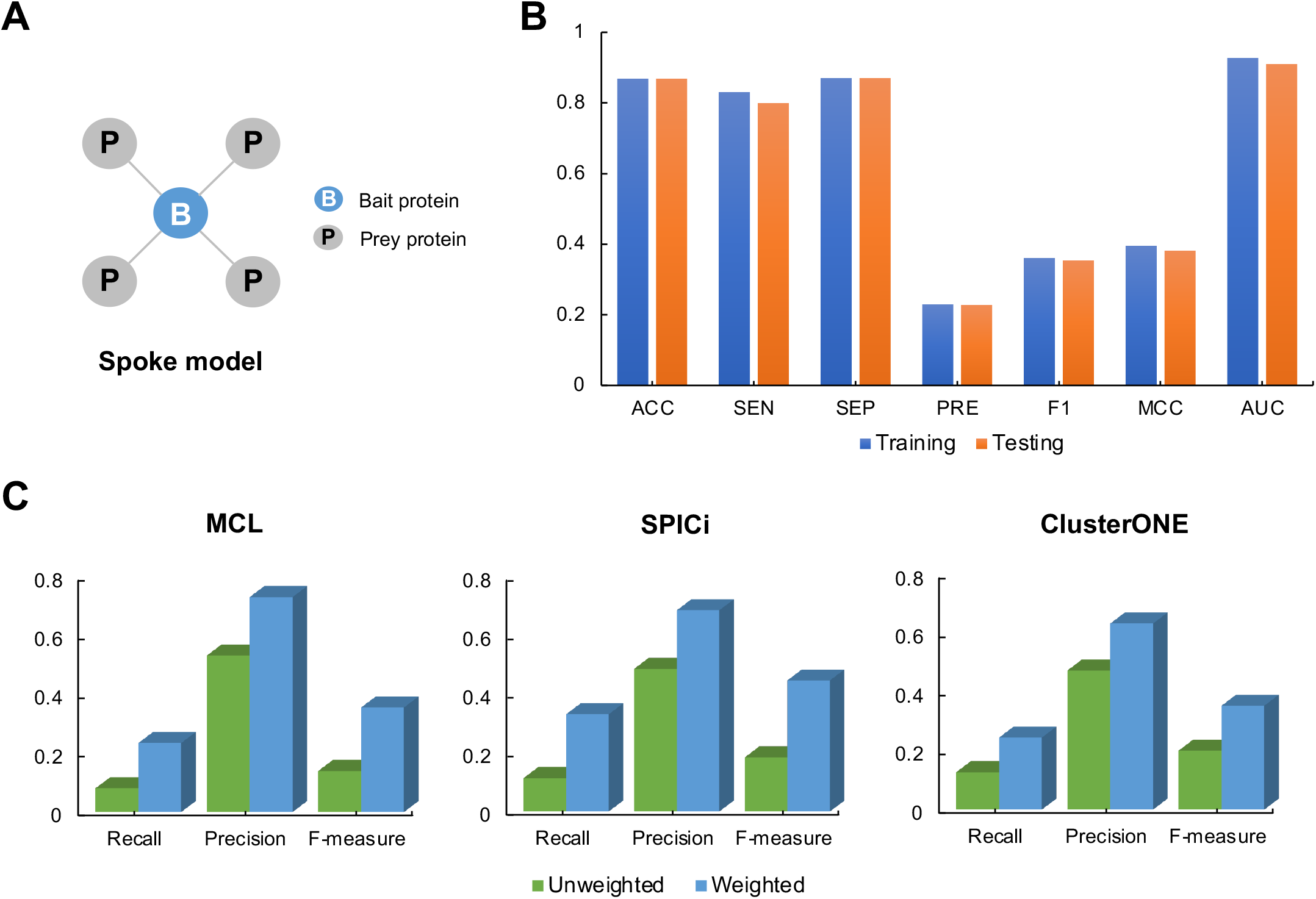
Building an accurate LE-PIN that is helpful for complex identification. **A**. In the spoke model, interactions are detected only between bait and prey proteins. **B**. Performance comparison of edge types classification on the training set and testing set. **C**. Performance comparison of three existing methods (MCL [8], SPICi [30] and ClusterONE [7]) on the weighted and unweighted PIN.

### Building a comprehensive atlas of human protein complexes

There are four major steps to build the atlas of human protein complexes. Firstly, constructing a PIN of featured edges (FE-PIN) based on the integrated PIN described above. Secondly, classifying the edges of FE-PIN to construct an PIN of labeled edges (LE-PIN). Thirdly, developing an effective method to accurately identify protein complexes from the LE-PIN of the test set. Lastly, applying the method to identify complexes from the LE-PIN of the integrated PIN and building a comprehensive atlas of human protein complexes.

#### Constructing the FE-PIN

We calculated features for each edge (or PPI) in the integrated PIN above to get a new PIN of featured edges, i.e., FE-PIN. Several different types of features were evaluated, including subcellular localization, PSSM-based features, GO semantic similarity, protein chain length and protein domain interactions. Eventually, 432 features in total were generated. Thus, each edge corresponds to a 432-dimensional feature vector.

#### Building the LE-PIN

Here, we built the LE-PIN of the integrated PIN in two steps. First, we trained a deep forest classifier [23] to label the edges of FE-PIN. We classified the edges in FE-PIN into two classes: c-edges that lie within complexes and nc-edges that lie outside complexes (see Materials and methods). The classifier outputs a probability as the weight for each edge. A larger weight indicates that the edge is more likely to be a c-edge, while a smaller weight indicates that the edge is more likely to be an nc-edge. To train and test the classifier, we generated the training and test sets by using complexes in the CORUM set. The deep forest classifier was evaluated using 10-fold cross-validation (Figure 2B). We can see that the performance results on the training set and the test set are quite similar in terms of different performance metrics (e.g., F1, MCC, and AUC), which indicates that the classifier is not over-fitting. Concretely, 80% of the c-edges (SEN = 0.800) and 87.2% of the nc-edges (SPE = 0.872) are correctly predicted on the testing set. To further verify that the weights of LE-PIN are helpful for complex identification, we applied three existing complex identification algorithms to the LE-PIN of the test set (Figure 2C). Here, we consider two cases: (a) the test set is treated as a normal PIN where all edges are weighted equally, so we also call it an unweighted PIN. (b) The test set is treated as a LE-PIN. Each method uses the default parameters, and the gold standard complex set is CORUM 2.0. We found that the performance on the weighted PIN (i.e., LE-PIN of the test set) is significantly better than that on the unweighted PIN, indicating that the weights in the LE-PIN are helpful to complex identification. With the classifier, we then labeled the unlabeled edges in FE-PIN, and got the LE-PIN.

#### Developing a novel and effective protein complex identification method

We developed a novel and effective method to accurately identify protein complexes from the LE-PIN. It is well known that protein interactions within complexes are strong and dense, while interactions between proteins of different complexes are weak and sparse. Thus, PPIs within complexes should correspond to c-edges of large weights, and PPIs between different complexes should correspond to nc-edges of small weights. We therefore designed an algorithm with an objective function aiming at high intra-complex cohesion based on c-edges and low inter-complex coupling based on nc-edges (see Materials and methods). **Table 1** shows the performance of different complex identification methods on the test set. Overall, our proposed method is superior to all the 15 state-of-the-art protein complex identification methods in terms of Recall, F-measure, MMR, and ACC, and performs comparably to the MCL method in terms of Precision. Although the MCL method has a higher Precision, its Recall is much lower, indicating that the predicted complexes matched only a small number of gold standard complexes, which consequently leads to a lower F-measure. F-measure considers both precision and recall and therefore reflects the prediction performance in a more balanced way. In summary, among all these methods, our method not only has good prediction performance (i.e., higher F-measure, MMR, and ACC) but also achieves the best balance between Recall and Precision, which shows that there is a better match between the complexes predicted by our method and the complexes of the gold standard set.

**Table 1.** Performance comparison between our method and existing methods on the testing set.

| Method | No. of total complexes | No. of small complexes | No. of large complexes | Recall | Precision | F-measure | MMR | ACC | Ref |
| --- | --- | --- | --- | --- | --- | --- | --- | --- | --- |
| MCL | 157 | 71 | 86 | 0.223 | **0.732** | 0.342 | 0.045 | 0.268 | [8] |
| EWCA | 1150 | 23 | 1127 | 0.126 | 0.257 | 0.169 | 0.031 | 0.207 | [28] |
| CFinder | 2071 | 1205 | 866 | 0.282 | 0.350 | 0.312 | 0.106 | 0.208 | [15] |
| GraphEntropy | 260 | 178 | 82 | 0.117 | 0.488 | 0.189 | 0.027 | 0.183 | [29] |
| ClusterONE | 243 | 28 | 215 | 0.223 | 0.634 | 0.330 | 0.048 | 0.254 | [7] |
| SPICi | 252 | 144 | 108 | 0.302 | 0.687 | 0.420 | 0.060 | 0.265 | [30] |
| MCODE | 25 | 11 | 14 | 0.010 | 0.320 | 0.020 | 0.003 | 0.116 | [9] |
| PC2P | 156 | 80 | 76 | 0.164 | 0.615 | 0.260 | 0.030 | 0.214 | [16] |
| IPCA | 763 | 116 | 647 | 0.191 | 0.413 | 0.261 | 0.041 | 0.188 | [10] |
| CORE | 163 | 62 | 101 | 0.104 | 0.436 | 0.167 | 0.021 | 0.179 | [11] |
| COACH | 463 | 5 | 458 | 0.107 | 0.184 | 0.135 | 0.018 | 0.187 | [12] |
| CMC | 6524 | 417 | 6107 | 0.393 | 0.157 | 0.224 | 0.105 | 0.205 | [13] |
| ProRank+ | 201 | 20 | 181 | 0.092 | 0.413 | 0.150 | 0.019 | 0.158 | [31] |
| DPClus | 167 | 20 | 147 | 0.123 | 0.455 | 0.193 | 0.023 | 0.210 | [14] |
| Clique | 51,112 | 1371 | 49,741 | 0.412 | 0.107 | 0.170 | 0.136 | 0.235 | [32] |
| HPC-Atlas | 999 | 841 | 158 | **0.608** | 0.702 | **0.651** | **0.178** | **0.273** |  |

*Note:* Five performance metrics are used, and for fair performance comparison, each method uses the default parameters. Bold number denotes the maximum value of this column. Ref denotes reference.

According to previous studies [33], a complex composed of two or three proteins was regarded as a small complex, and those contain more than three proteins were treated as large complexes. There are much more small complexes than large complexes in the CORUM set, which indicates that small complexes may dominate large complexes in human cells (e.g., small complexes account for 65% of the CORUM 3.0). Many existing identification methods of protein complexes are more effective in identifying large complexes because they take densely connected subnetworks of PPI networks as complexes. Therefore, small complexes composed of single or two edge are easy to be missed by those methods. Our method can find more small complexes than most existing methods.

#### Building a comprehensive atlas of human protein complexes

Having created the accurate LE-PIN and an effective complex identification method, we next used the method to identify complexes from the LE-PIN of the integrated PIN and build a comprehensive atlas of human protein complexes. In total, 8944 complexes were eventually identified, which were used to construct our atlas of human protein complexes.

### Identifying previously uncharacterized protein complexes

To assess HPC-Atlas’s ability to identify novel protein complexes, we first downloaded the latest CORUM (version 3.0) and generated an independent set of protein complexes that are not included in the previous CORUM set used for training and testing. **Table 2** presents the number of protein complexes predicted by different methods on the LE-PIN of the integrated PIN. The parameters of each method use the default parameters. Five of the fifteen methods failed to work on large PPI networks, so they are not included in Table 2. We found that our method got the largest number of successfully matched complexes, up to 272, of which 12 are exactly matched with the complexes in the independent set. Here, a successful match means that the matching rate (defined in Equation (17)) of a predicted complex with a certain true complex is no less than 0.2, and an exact match means that the matching rate is 1.0. The numbers of complexes successfully matched of existing methods are much smaller than that of our method. Interestingly, the scores (defined in Equation (28)) of the 12 exactly matched complexes identified by our method are about 0.5. Therefore, in the sequel we regard the complexes with scores no less than 0.5 as real complexes, because they have stronger cohesion and lower coupling. Similarly, we found that small complexes account for the majority of the complexes identified by our method.

**Table 2.**
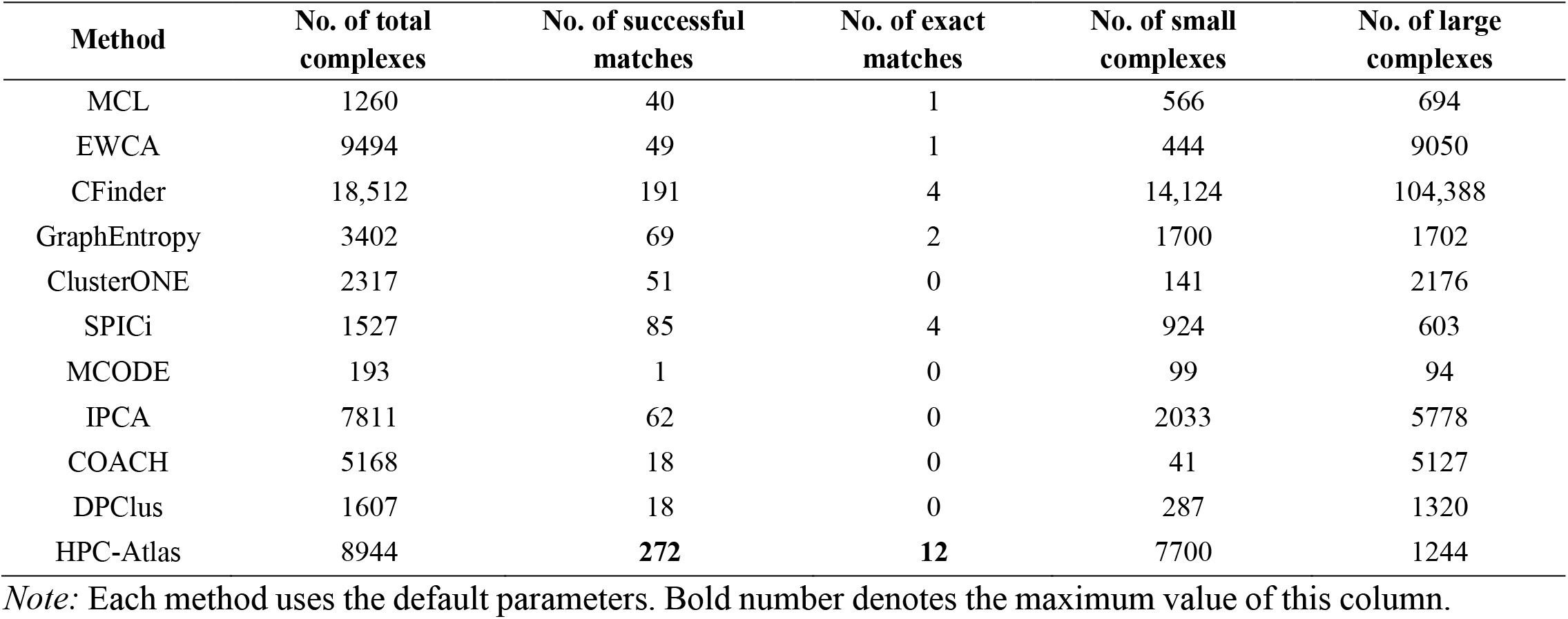
Comparison of the numbers of complexes identified by different methods from LE-PIN.

As an example of the effectiveness of HPC-Atlas in identifying new protein complexes, **Figure 3**A, 3B show two protein complexes predicted by our method, one of which is a new complex (Figure 3B) and the other is a complex that has an exact match in the independent set (Figure 3A). As shown in Figure 3A, the EIF2B complex comprises five proteins that belong to a family of EIF2B involving in catalyzing the exchange of eukaryotic initiation factor 2-bound GDP for GTP. Our method assigned a score of 0.509 to the EIF2B complex whose most significantly enriched functional annotations include translation initiation factor activity and eukaryotic translation initiation factor 2B complex, which are biologically consistent with EIF2B’s functions. However, the other methods cannot identify this complex. Another complex is denoted by UGT2 (Figure 3B), because all its proteins belong to the UDP-glycosyltransferase family and have the function of eliminating potentially toxic xenobiotics and endogenous compounds. UGT2 has a high score, but is not in the latest CORUM set. With a further analysis, we found that proteins in this complex have quite similar subcellular locations, especially UGT2A1 has completely consistent subcellular locations with UGT2A3. This indicates that these proteins have high probability to form the complex to carry out essential functions of cells. Meanwhile, we noticed many significantly enriched functional annotations among the member proteins of this complex, including glucuronidation and metabolism of drugs and xenobiotics. In summary, it is likely that UGT2 is a real complex that participates in UDP-glucuronosyltransferases catalyze reaction, and eliminates toxic xenobiotics and endogenous compounds because it contains proteins of similar functions and locations in human cells. Overall, these results demonstrate that HPC-Atlas is able to identify potential novel protein complexes.

**Figure 3.**
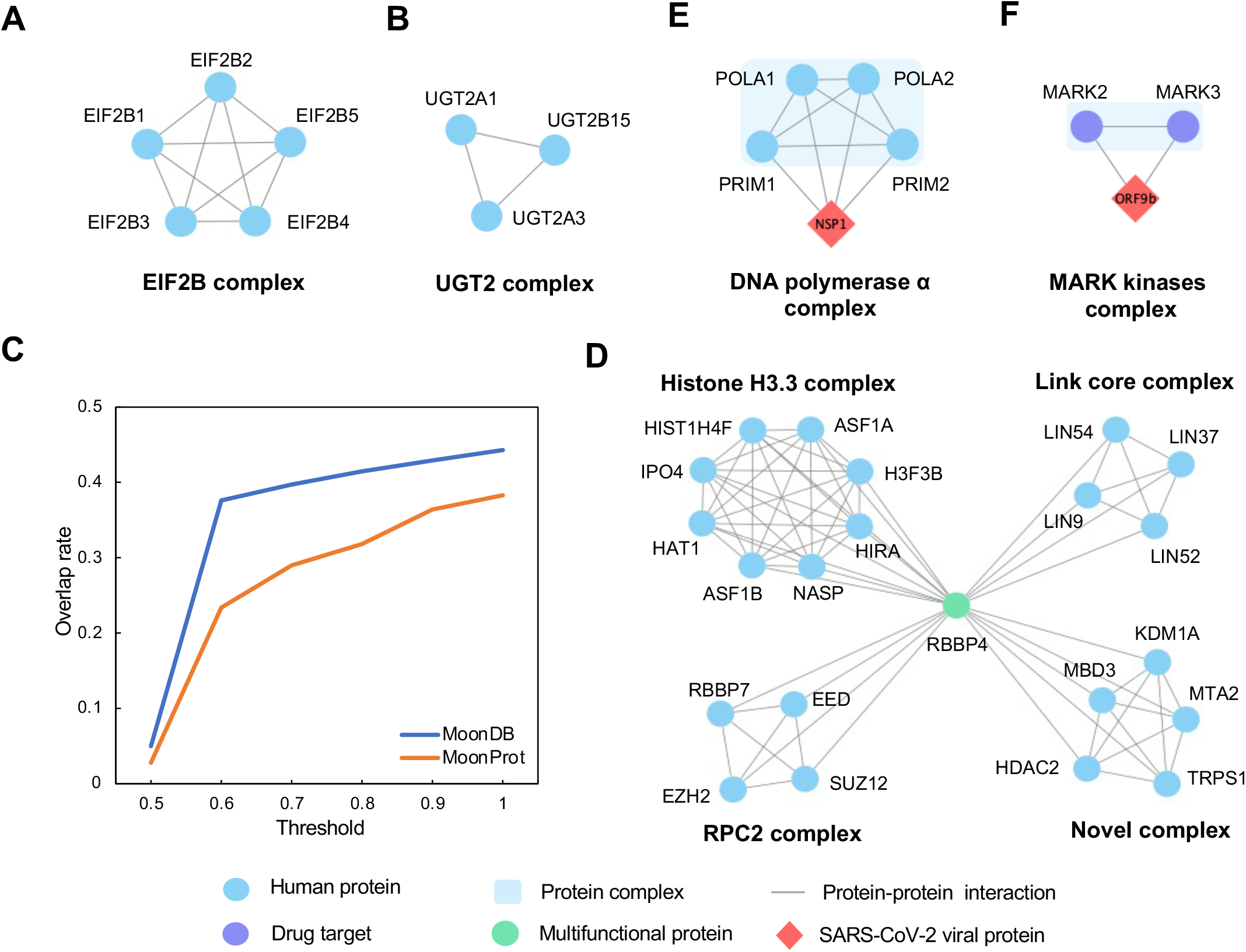
HPC-Atlas is useful in discovering new biological findings, including predicting novel complexes, multifunctional proteins and SARS-CoV-2 affected human protein complexes. **A**. The predicted EIF2B complex has an exact match in the independent set. **B**. Our method identified UGT2, a previously unreported complex. **C**. The overlap ratios between the multifunctional proteins we identified and these in MoonDB [36] and MoonProt [37] databases at different complex overlap thresholds. **D**. Multifunctional protein RBBP4 participates in four distinct complexes, three of which are included in the CORUM set and the rest one is a new complex. **E**. Example of SARS-CoV-2 NSP1 interactions with human proteins, which form the DNA polymerase alpha complex. **F**. The MARK kinases complex in the HPC-Atlas contains two drug targets, both of which interact with SARS-CoV-2 ORF9b. SARS-CoV-2, Severe acute respiratory syndrome coronavirus 2.

### Identifying multifunctional proteins

Multifunctional proteins, also known as multitasking proteins, are a type of proteins that perform multiple biochemical functions [34]. It has been found that multifunctional proteins are associated with human diseases and targets of drugs [34]. However, it is not clear how many multifunctional proteins there are in human, and there are few effective methods for identifying multifunctional proteins [35]. Two existing databases [36,37] contain only a small number of human multifunctional proteins, therefore they cannot completely cover human multifunctional proteins. One of the databases, MoonProt [37], contains 107 moonlighting proteins (a subclass of multifunctional proteins), which execute unrelated biological functions. The other database, MoonDB [36], contains 282 human proteins, including moonlighting and multifunctional proteins. Now, as we have a comprehensive atlas of human protein complexes, from which we can potentially identify multifunctional proteins.

According to a recent work [17], multifunctional proteins participate in multiple distinct complexes and perform two or more biological functions. Therefore, we first constructed a set of protein complexes with limited overlapping, by removing the complexes that have high overlap with the other complexes (see Materials and methods), and then treated the proteins appearing in at least two complexes in the set as multifunctional proteins. Thus, we identified a total of 456 multifunctional proteins, which make up 6% proteins in the set of complexes. Compared with the 364 known multifunctional proteins in the existing databases (i.e., MoonProt and MoonDB), there is a significant lift in coverage of human multifunctional proteins. We found that these multifunctional proteins usually participate in 2−4 complexes, and most of them are involved only in two complexes. Additionally, we compared the overlap between the multifunctional proteins in these two databases and the multifunctional proteins obtained by our method under different complex overlap thresholds (defined in Equation (32)). As show in Figure 3C, with the increase of the threshold, the overlap ratio of multifunctional proteins significantly increases. This suggests that a lower threshold can get more robust results and identify more novel multifunctional proteins. Meanwhile, this also shows the potential of our atlas to identify moonlighting proteins, which are the most essential and fascinating proteins in multifunctional proteins.

Figure 3D shows an example of multifunctional protein, RBBP4, participating in four protein complexes. Among these complexes, three of them are included in the latest version of the CORUM set, and the other is a novel complex that has not been reported. According to the description of complexes in the CORUM set, we know that the PRC2 complex maintains the transcriptionally repressive state of many genes, the Histone H3.3 complex mediates DNA-synthesis by containing distinct histone chaperones, and the Link core complex plays an important role in cell cycle-dependent activation. We further analyzed the potential functions of RBBP4 in these complexes using Reactome annotation enrichment. The enriched annotations of RBBP4 in PRC2, Histone H3.3, Link core and the novel complex are transcriptional regulation by E2F6, cellular senescence, cell cycle, and regulation of PTEN gene transcription, respectively, which are known functions of RBBP4. We therefore believe that RBBP4 is involved in the four complexes of different functions, which is likely to be a multifunctional protein. This example illustrates the potential of our atlas to identify multifunctional proteins from the constructed set of complexes of limited overlap.

### SARS-CoV-2 affected human protein complexes

Severe acute respiratory syndrome coronavirus 2 (SARS-CoV-2), as the causative agent of coronavirus disease-2019 (COVID-19), has led to more than 660 million confirmed cases and 6.6 million deaths globally, and seriously affected people’s normal life [38]. The SARS-CoV-2 proteins potentially interact with multiple human proteins after the virus infects human cells [39]. These interactions involve several complexes and biological processes, including DNA replication, vesicle trafficking, and lipid modification [39]. Therefore, identifying protein complexes that may be affected by the virus will help us to develop drugs against SARS-CoV-2. Fortunately, a recent study [39] has identified 332 high-confidence PPIs between human and SARS-CoV-2 proteins and 66 druggable human proteins through AP-MS experiments. Here, we tried to find SARS-CoV-2 relevant protein complexes with our HPC-Atlas to further demonstrate its value in exploring biological findings.

From HPC-Atlas, we identified 751 human protein complexes that contain at least one protein interacting with a SARS-CoV-2 protein. We call them SARS-CoV-2 affected complexes. These complexes contain virus interacting proteins ranging from one to seven, and most of them contain only one or two such proteins. Among them, all the proteins contained in the 21 complexes associate with the SARS-CoV-2 proteins. Using the latest version of the CORUM set as the gold standard, we found that 296 complexes (nearly 40%) have matches in the CORUM set (version 3.0), of which 17 complexes have exact matches. For example, DNA polymerases are a group of polymerases that are often used in amplification techniques (e.g., PCR amplification and loop-mediated isothermal amplification) to detect the presence of SARS-CoV-2 sequences [40,41]. Here, we identified a DNA polymerase alpha-primase complex that contains four proteins, all of which interact with the NSP1 protein of the SARS-Cov-2 viral (Figure 3E). We then performed functional enrichment analysis on the complex and found that it has strong enrichment of GO terms specific to DNA replication and synthesis of RNA primer, and is highly specific for the nasopharynx tissue. These are consistent with the characteristics of DNA polymerases.

Additionally, we found 143 of the above complexes contain drug target proteins, suggesting that these complexes may contribute to drug development. For example, we identified MARK kinases complex that contains two proteins, which are all drug targets interacting with the ORF9b protein of the SARS-CoV-2 viral (Figure 3F). MARK2 and MARK3 have been approved as drug targets for the treatment of cancer and myelofibrosis [42]. Furthermore, we found high overlap between the subcellular localizations of MARK2 and MARK3, indicating that they may form a complex. Recent studies have also shown that drug repurposing is a promising scheme for exploring potential SARS-CoV-2 drug targets [42,43]. We therefore believe that this complex might contribute to fighting COVID-19.

## Discussion

A comprehensive atlas of human protein complexes can help us to better understand the critical functions and mechanisms in human cells, and find the related biological pathways of diseases. However, our ability to interpret unknown biological problems is limited by the lack of a reliable atlas of human protein complexes. Herein, we present HPC-Atlas, the most accurate and comprehensive atlas of human protein complexes to date, which expands our knowledge about protein complexes in human cells.

### Building a more comprehensive protein interaction network

Since high-throughput techniques usually miss some interactions (e.g., prey-prey interactions) and introducing false positives, we used the L3 and WMM algorithms to identify high confidence false negative PPIs from HuRI and BioPlex respectively. Then, integrating the expanded networks to increase the coverage of the human interactome. Five categories of features were evaluated to represent the interactions. This allows our deep forest classifier to accurately predict labels for protein pairs. We finally generated a high-quality PIN (we called LE-PIN) where edges are weighted with probabilities predicted by the classifier, which indicate whether the edges fall in or outside complexes.

### Accurately identifying more protein complexes

In the past decade, many computational methods have been proposed to identify complexes from PINs. Most of them used clustering algorithms, which regard protein complexes as dense subgraphs of the PIN. However, they did not differentiate the PPIs (or edges in PINs) except for assigning them different weights or confidences. In this paper, we classified the edges into complex edges (i.e., c-edges) and non-complex edges (i.e., nc-edge), and regard complexes as subnetworks of proteins connected by c-edges. This is a novel idea about protein complexes. Based on this idea, we developed a novel algorithm to accurately identify complexes based on the LE-PIN. Our method considers that a complex is a subgraph of c-edges and with high cohesion and low coupling, it iteratively adds and removes proteins to maximize the complex’s score. Performance evaluation shows that our method substantially outperforms existing methods, especially in terms of Recall and F-measure on the testing set. To illustrate that our method can effectively identify novel complexes, we generated an independent test set that does not overlap with the CORUM set (version 2.0). The results show that our method can predict more complexes with higher accuracy. In addition, a number of novel complexes with high score were verified by significantly enriched functional annotations.

### Exploring new biological findings

As a valuable resource, HPC-Atlas can be used in many fields, including structural biology, systems biology, and disease-related molecular biology. In this work, to illustrate that the atlas can be used as a new biological discovery source, on the one hand, we used the atlas to identify multifunctional proteins. Currently, there are few effective methods of multifunctional protein identification. With our atlas, we can search such proteins from a set of complexes with limited overlap. We also found that our atlas has the potential of identifying moonlighting proteins, a special sub-class of multifunctional proteins, by searching the existing databases. On the other hand, we used the atlas to find SARS-CoV-2 affected complexes. We can identify the affected complexes that contain at least one protein interacting with a SARS-CoV-2 protein. Certainly, we can also use the atlas to identify other disease-related complexes.

Taken together, HPC-Atlas is not only an extensible computing framework that can integrate additional PPI sources to identify complexes, but also a valuable resource for biological knowledge discovery. In the future, we plan to combine more PPI data and construct a more comprehensive map to further boost the prediction performance of complexes, and apply the map to other biomedical problems such as new drug discovery for emerging diseases.

## Materials and methods

### Datasets

#### Protein interaction datasets

We used two latest protein interaction datasets or networks HuRI and BioPlex as the basic data. The HuRI network derived from Y2H experiments, consists of 63,132 PPIs involving 8985 proteins. The BioPlex network was the results of AP-MS experiments from 239T and HCT116 cells, comprising 167,932 PPIs across 14,484 proteins.

With these two PINs above, an expanded and integrated PIN was generated. First, we applied the L3 and WMM methods, whose effectiveness has been shown in previous studies [17,24,26], to identify new high-confidence interactions from HuRI and BioPlex, respectively. WMM is based on the hypergeometric distribution (Equation (1)), which can be used to calculate the probability of PPIs in PIN. For a given PIN, the probability of interaction between two proteins A and B is:

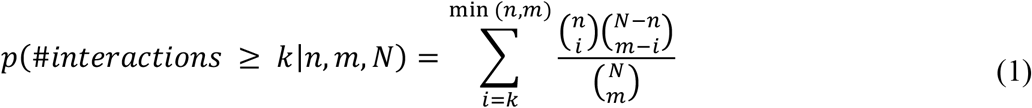

where *k* is the number of interactions between two proteins A and B, *n* and *m* are the total number of interactions for protein A and protein B, respectively. *N* represents the total number of interactions in the PIN. WMM calculates a p-value for each pair of proteins to represent the confidence of the corresponding interaction and rank the interactions according to their p-values. We then selected these PPIs with p-value less than a certain threshold. Here, the threshold is the minimum confidence value (i.e., the larger p-value) obtained by matching the predicted PPIs and the real PPIs in the BioPlex. When applying L3 to the HuRI network, we considered only the top 6000 predicted interactions as true PPIs following previous research [24]. L3 adds edges based on the principle of complementary interfaces (see Figure S2). The authors of L3 believed that two interacting proteins often have complementary interfaces. Therefore, two proteins X and Y with similar interface usually have many common neighbor proteins. And the neighbor proteins of X may interact with protein Y because they have complementary interfaces, and this interaction can be detected by using paths of length 3 (L3). We then integrated these two PINs after adding the predicted new interactions to construct an integrated PIN, which contains 2,658,160 PPIs over 16,632 human proteins. The statistics of PINs in this study are shown in Table S1.

#### The training and testing sets for classifying edges

To train and evaluate the deep forest classifier, we created a training set and a testing set based on the CORUM core set (version 2.0). The CORUM set was randomly divided into training and testing sets. Complexes in each set were disassembled into protein pairs and removed redundant protein pairs. A pair of proteins is defined as a c-edges if it falls at least in a complex, and as an nc-edges if the two proteins do not fall in any complexes. The final training and testing sets were generated by mapping the training and testing complexes from CORUM 2.0 into the integrated PIN, which are comprised of 4012 and 3513 c-edges, 87,050 and 74,490 nc-edges, respectively.

#### Gold standard protein complex sets

To evaluate the complex identification method on the LE-PIN of the testing set, we used CORUM 2.0 and filtered out all non-human complexes. And we removed the complexes with sizes less than 2 from the CORUM set. Additionally, we also downloaded the latest CORUM set (version 3.0) and generated an independent set of protein complexes that were not included in the CORUM 2.0. This independent set contains 518 human protein complexes, which were used to evaluate our method and to compare it with existing ones in identifying new complexes. Table S2 summarizes the protein complex sets used in this study.

### Calculating protein interaction features

To characterize the PPIs from different aspects, we generated 432 features for each interaction or protein pair, which can be divided into five groups: subcellular localization, PSSM-based features, gene ontology semantic similarity, protein chain length and protein domain interactions. All features were standardized by scaling them to the range [0,1].

#### Subcellular localization

Protein subcellular localization provides information about the specific location of a certain protein in human cells, such as nuclear region, cytoplasm and cell membrane. Therefore, a pair of interacting proteins should be close to each other, and a complex should contain proteins that are also close to each other to perform related biological functions in cells. In this work, we calculated four location-based features using the subcellular localization annotations of each protein downloaded from the UniProt database [44]. Specifically, let 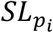 and 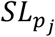 represent the subcellular localization annotation sets of *protein*_*i*_ and *protein*_*j*_ (*i* ≠ *j*). The following four location-based features for interacting proteins *p*_i_ and *p*_j_ are evaluated:

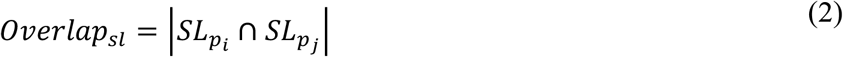

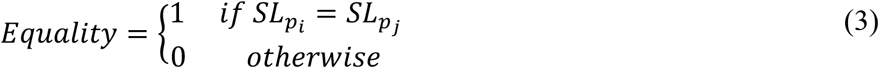

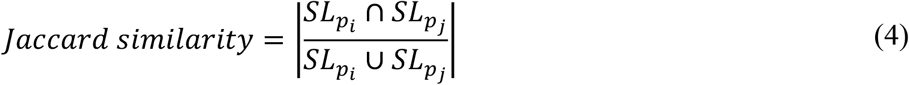

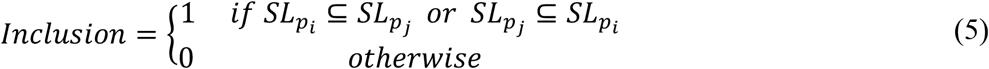

#### Position-specific scoring matrix (PSSM) based features

The evolutionary conservation information of PSSM derived from protein sequences is widely used. The PSSM-based features have been successfully used to promote the performance of protein property prediction [45−48]. Here, we used the MEDP [49] method to generate a 420-dimensional feature vector by using the PSSM file of each protein as input. The PSSM of a protein was evaluated by multiple sequence alignments with PSI-BLAST [50] to search against NCBI non-redundant database [50]. Then, the PSSM-based features of a protein pair were composed of the sum of feature vectors of the two proteins and the similarity of the two feature vectors calculated by Euclidean distance.

#### Gene ontology semantic similarity

Gene ontology (GO), as a widely used biological data resource, provides a convenient way to evaluate the semantic similarity of pairwise GO terms [51]. Inferring semantic similarity between GO terms has been successfully used in many research areas, such as protein function prediction and gene network analysis [52]. The semantic similarity of GO was calculated by using the structural relationships, including parent-child relationships and sibling relationships among nodes in the ontology. A protein usually contains at least one GO term. In this work, we use the GOSemSim package [53] to calculate GO semantic similarity between a pair of proteins, and get a similarity score as the final feature value.

#### Protein chain length

Proteins have a variety of conformations, and their chain lengths are generally between 50 and 2000 amino acids [54]. Large proteins are usually composed of several different protein domains and structural units [54]. Therefore, we want to know whether their protein chain lengths of interacting proteins are similar. The absolute difference between the chain lengths of a pair of proteins was used as the feature value. The length of each protein chain is obtained from the UniProt database [44].

#### Protein domain interactions

Protein domains are a class of structural units in proteins [55]. The interaction between a pair of proteins is usually related to the physical interactions between their specific domains [56]. Protein domains are essential to understand protein interactions and enable us to have a more comprehensive understanding of protein functions and PPI networks. Protein domain interactions and each protein’s domain annotations were downloaded from Pfam [57] and Uniprot databases, respectively. Concretely, let 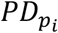 and 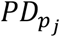 be the sets of protein domains of a protein pair (*p*_*i*_, *p*_*j*_)_*i*≠*j*_, *PD_Pfam* is a set of protein domain interactions. 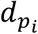and 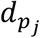are interacting domains from 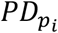 and 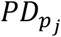 respectively. Then, the following protein domain interaction features are defined:

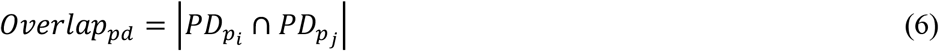

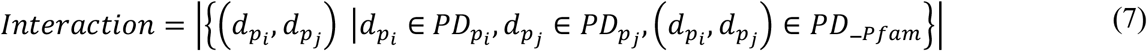

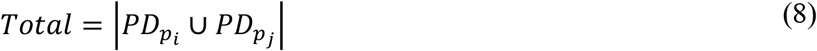

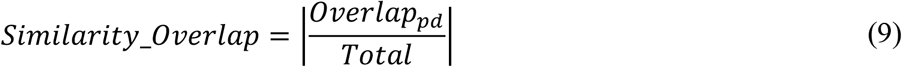

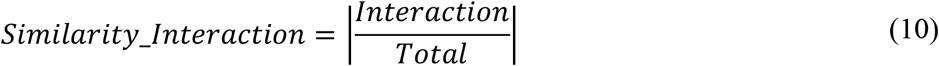

### Evaluation metrics

The evaluation of our method consists of two parts: edge labeling and protein complex identification, which correspond to a classification task and a clustering task, respectively.

#### Performance evaluation of edge labeling

The performance of the deep forest classifier for edge labeling is evaluated through 10-fold cross-validation. Since there are more nc-edges than c-edges in the training set, traditional cross-validation is easily dominated by the nc-edges, which results in prediction bias. To handle the imbalanced data set, we performed 10-fold cross-validation by using the sub-sampling strategy. First, the c-edges and nc-edges in the training set were randomly split into ten subsets, respectively. In each round, nc-edges were randomly selected from nine subsets of nc-edges, combined with nine subsets of c-edges to create a 1:1 balanced training set, while the remaining edges were merged as the testing set. For a comprehensive assessment of the classifier, we used seven performance measures, including accuracy (ACC), sensitivity (SEN), specificity (SEP), precision (PRE), F1-score (F1), Matthew’s correlation coefficient (MCC) and the area under the receiver operating characteristic curve (AUC), which are evaluated as follows:

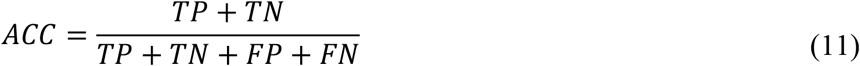

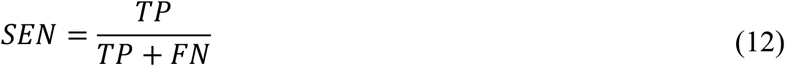

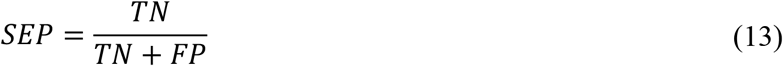

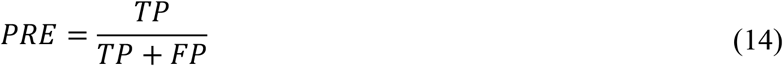

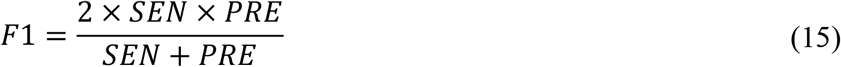

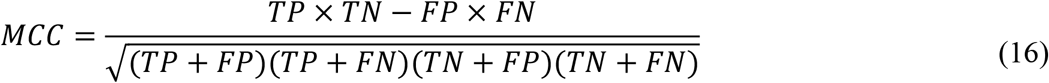

where *TP, FP, TN, FN* represent the numbers of true positives (i.e., correctly predicted c-edges), false positives, true negatives and false negatives, respectively.

#### Performance evaluation of protein complex identification

To evaluate protein complex identification, we first check whether a predicted complex matches some complexes in the gold standard complex set, i.e., the CORUM set (version 2.0 or 3.0). Here, the matching rate between a predicted complex and a gold standard complex is calculated as follows:

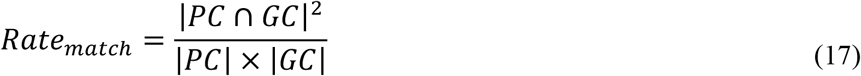

Above, *PC* and *GC* represent the numbers of proteins contained in the predicted complex and the gold standard complex, respectively. If *Rate*_*match*_ ≥ 0.2, we regard that *PC* and *GC* match successfully, which is consistent with the previous research definition [28,58–60]. Five widely-used metrics are adopted to evaluate the proposed method, including Recall, Precision, F-measure, maximum matching ratio (MMR), and geometric accuracy (ACC), which are calculated as follows:

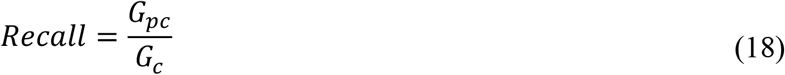

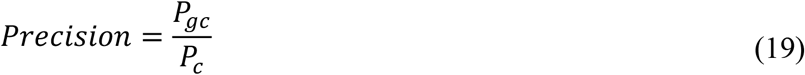

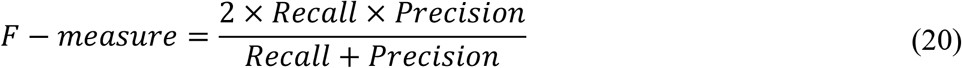

where *G*_*c*_ is the number of complexes in the gold standard set and *G*_*pc*_ is the number of gold standard complexes matched with some predicted complexes, *P*_*c*_ is the number of predicted complexes and *P*_*gc*_ is the number of predicted complexes matched with some gold standard complexes. F-measure is the harmonic mean of Recall and Precision.

MMR [7] is defined as the sum of the maximum matching edge weights between the gold standard set and the predicted complex set divided by the number of gold standard complexes. The maximum matching edge is obtained by building the maximal matching in a bipartite graph between the gold standard and predicted complexes, and the edge weight is given by the *Rate*_*match*_.

ACC [61] is obtained by computing the geometrical mean of sensitivity (SN) and positive predictive value (PPV). Let *t*_*g,p*_ be the number of shared proteins between the gold standard complex *g* and the predicted complex *p*, and *N*_*g*_ be the number of proteins in the gold standard complex *g*.

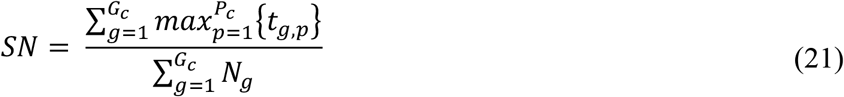

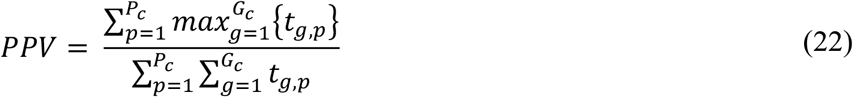

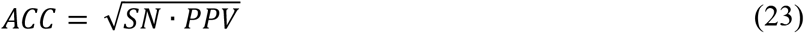

### Classifying the edges of FE-PIN

We used the deep forest [23] algorithm called gcForest to classify the edges of FE-PIN into two types: c-edges and nc-edges. gcForest simulates the hierarchical structure of a neural network. Each layer uses a group of forests, and the forests of each layer use the information of the previous layer as input and their output is taken as input of the next layer. Comparing with other deep learning methods, gcForest has much fewer hyper-parameters and is not limited to the size of training data. That is, it also shows good performance on small data sets. We therefore used gcForest to classify edge types in FE-PIN. To boost performance, we also concatenated the lightgbm [62] predictor with gcForest. We used the default parameters of the gcForest algorithm. We trained the classifier by using 10-fold cross-validation on the training set and then evaluated the classifier on the testing set. The classifier’s output probability for each edge is taken as its weight in the LE-PIN. With all edges of FE-PIN labeled or weighted, we got the LE-PIN.

### Identifying protein complexes

We designed a novel approach to identify protein complexes from the LE-PIN generated by the edge classification step described above. First, we represented the PPI network as a weighted undirected graph *G* = (*V, E, W*), where *V* represents the set of nodes (i.e., proteins) in the network, *E* is the set of edges, each of which corresponds to interacting protein pairs, *W* represents the weights on the edges, each of which is the probability value predicted by the deep forest classifier. We took these edges with weight greater than 0.5 as c-edges (i.e., the corresponding pairs of proteins lie in a complex), and the rest as nc-edges. Then, we defined a complex scoring function, which consists of two parts: density and modularity. Density represents the cohesion of the subgraph (SG), and modularity represents the coupling degree of the subgraph. The density *D*_*SG*_ of the subgraph *SG* is evaluated as follows:

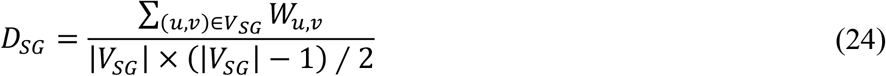

where *u* and *v* are two nodes within the subgraph, 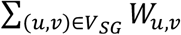 is the sum of internal edge weights of the subgraph, and *V*_*SG*_ is the node set inside the subgraph. The modularity *M*_*SG*_ of the subgraph *SG*, is defined as follows:

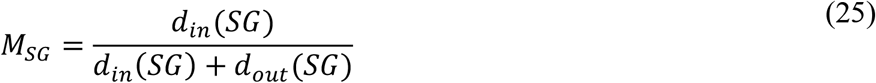

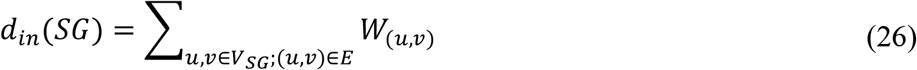

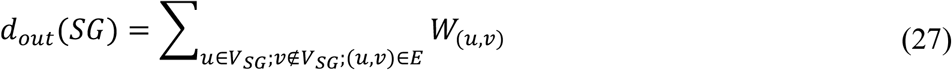

Above, *d*_*in*_(*SG*) is the sum of weights of all edges in the subgraph, and *d*_*out*_(*SG*) is the sum of weights of edges between the inner and neighbor nodes of the subgraph. If the subgraph has high modularity, it means that the subgraph is dense, but sparsely connected with outside nodes.

The score of a protein complex *F*_*SG*_ is evaluated by combining the density and modularity of the subgraph as follows:

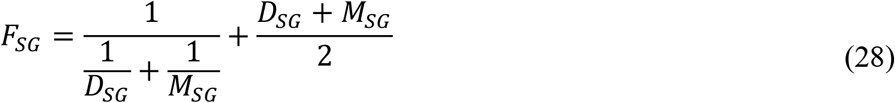

Generally, if a subgraph has a high *F*_*SG*_ score (i.e., the subgraph has a high density and modularity value), it is more possibly a protein complex. Our method identifies a protein complex by iteratively adding and removing nodes to maximize the *F*_*SG*_ score. In detail, we have four criteria for selecting the most appropriate node in each iteration from the neighbor node set of the subgraph to expand the subgraph:

(i) The node *n*_*s*_ should have edges connecting with the internal nodes of the subgraph, and their weights should be greater than 0.5.

(ii) The node *n*_*s*_ should satisfy Equation (29).

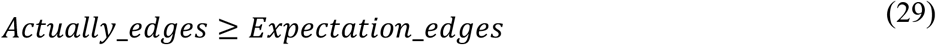

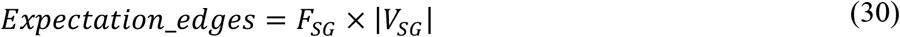

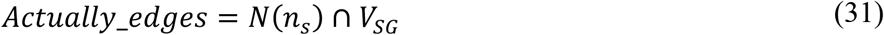

That is, *Actually_edges* is greater than *Expectation_edges*. In Equation (31), *N*(*n*_*ε*_) is the set of neighbor nodes of *n*_*ε*_.

(iv) The *F*_*SG*_ score should increase after adding a certain node.

(v) Selecting the node that can maximize *F*_*SG*_ from the neighbor nodes that meet the above three criteria to join the subgraph. At the same time, deleting the selected node from the neighbor node set. Iteratively adding nodes till the neighbor node set is empty or *F*_*SG*_ can increase no more.

When the process of adding nodes ends, we iteratively remove nodes from the subgraph to maximize *F*_*SG*_. Specifically, in each iteration we remove a node from the boundary node set of the subgraph to optimize the subgraph by following three criteria as follows:

(i) The *F*_*SG*_ score should increase after removing a certain node from the boundary node set of the subgraph. The boundary node set of the subgraph is a subset of the internal nodes of the subgraph, where each node has edges to connect both internal nodes and external nodes of the subgraph.

(ii) The node should satisfy Equation (29).

(iii) Selecting the node that can maximize *F*_*SG*_ from the boundary nodes that meet the above two criteria to remove from the subgraph.

The process of removing nodes from the subgraph continues till *F*_*SG*_ reaches stable, and the final complex is then output. Details of the process above are outlined in Algorithm 1.

In implementation, our method uses multithreading technology, and each thread handles several seed nodes. To speed up the calculation, we sorted the seed nodes by k-shell decomposition [63]. We found that k-shell decomposition is very suitable for sorting seed nodes in PIN generated by high-throughput techniques. This method can effectively overcome the bias of the spoke model. If the k-shell value of a protein is very large, it means that a large amount of computation is needed to form a complex based on the protein. Therefore, each thread handles either only one or two nodes with a high k-shell value or multiple nodes with a low k-shell value. In this work, the k-shell values ≥ 20 are considered as high values, and the rest are considered as low values, which can be set by the user in the algorithm. After the above complex search process is completed, we discard duplicate protein complexes from the generated candidate complex set to generate the final complex set. The duplicate protein complexes mean that the matching rate of the two complexes is 1.0.

#### Algorithm 1 Protein complex identification

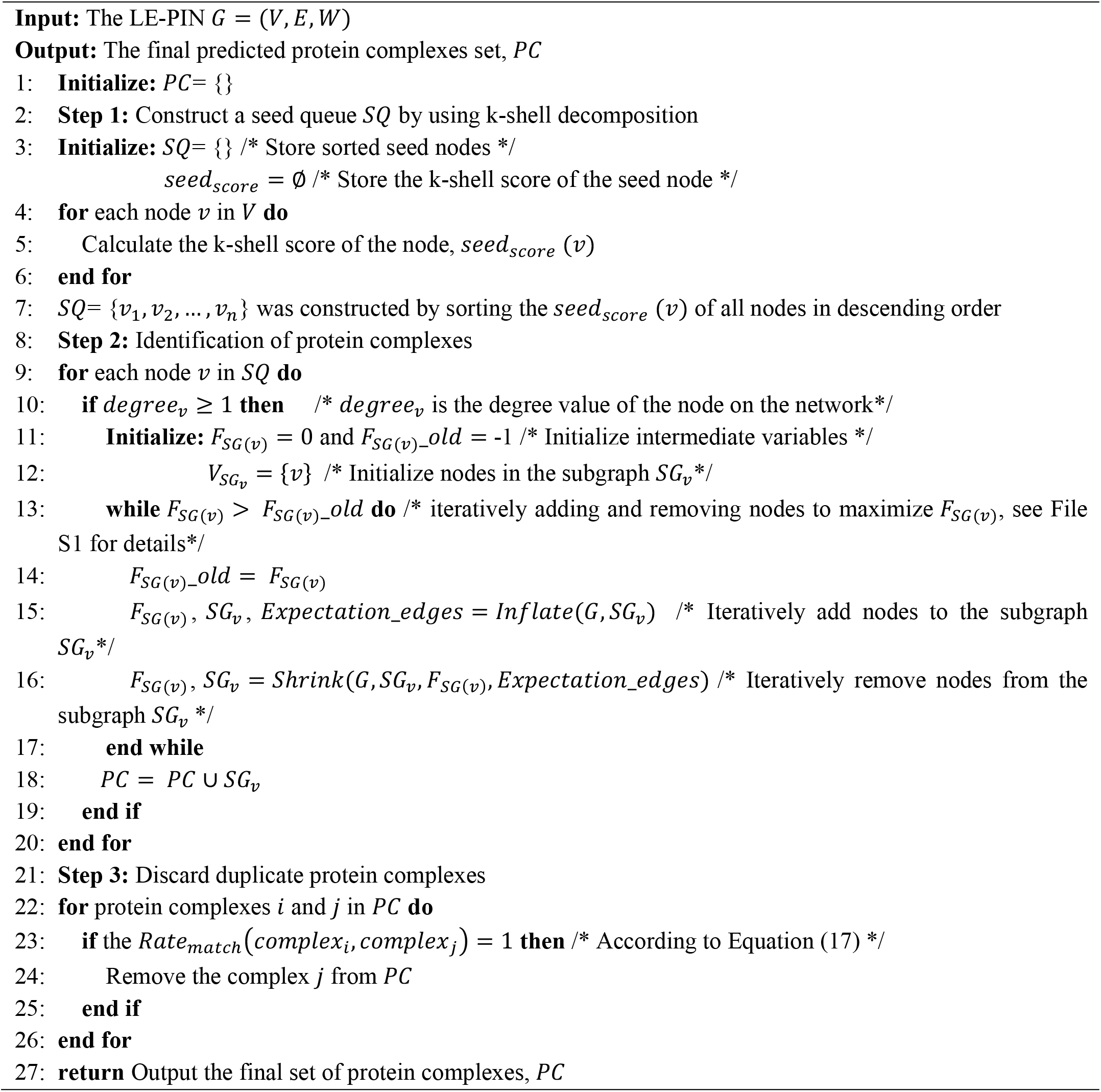

### Identifying multifunctional proteins

Multifunctional proteins are extremely important proteins, which play a central role in many diseases, but there are few effective methods to predict multifunctional proteins [35,37]. As previous studies [17], in order to identify multifunctional proteins, we first built a complex set with limited overlap. *SubComplex_index* was used to determine the degree of overlapping between two complexes as follows:

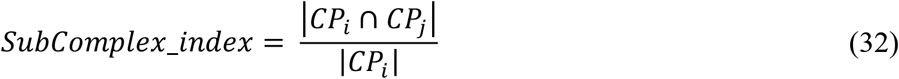

where *CP*_*i*_ and *CP*_*j*_ are the sets of proteins contained in the two complexes. We finally constructed a complex set by selecting the complexes with *SubComplex_index* < 0.5 from our final protein complex set (i.e., 8944 complexes). With this selected complex set, we then found these proteins that appear in two or more complexes, which were considered as multifunctional proteins.

### Identifying new protein complexes

To identify new complexes, we first rank the complexes identified from the LE-PIN according to their scores (defined in Equation (28)). The higher the complex’s score is, the more likely it is to be a real complex. Then, we exactly matched complexes in the independent set with all the identified complexes (i.e., 8944 complexes), and took the lowest score of all matched identified complexes as a threshold. With this threshold, we regard any identified complex with a score no less than the threshold as real complexes. In this paper, the score threshold is 0.5. Thus, we took these identified complexes having a score no less than 0.5 as real complexes. In addition, we also performed functional enrichment analysis for each complex with g:Profiler [64], which contains the most popular data sources: GO [65], Reactome [66], CORUM [6], KEGG [67], and Human Phenotype Ontology [68] (HPO). The g:Profiler method was usually used for enrichment analysis of complexes, i.e., detecting statistically significantly enriched biological processes by mapping user-provided protein lists to data sources. Here, we applied the g:SCS method to obtaining the p-value of each complex and ignored the electronic annotations. All proteins in the integrated PIN were used as the background set.

## Data availability

HPC-Atlas provides a user-friendly webserver at http://www.yulpan.top/HPC-Atlas.

## Code availability

The data set, method code and result file of HPC-Atlas can be downloaded at https://github.com/yul-pan/HPC-Atlas.

## CRediT author statement

**Yuliang Pan:** Conceptualization, Methodology, Software, Writing - original draft. **Ruiyi Li:** Methodology, Investigation. **Wengen Li:** Conceptualization, Validation. **Liuzhenghao Lv:** Software. **Jihong Guan:** Conceptualization, Supervision, Writing - review & editing, Funding acquisition. **Shuigeng Zhou:** Conceptualization, Methodology, Writing - review & editing, Supervision, Funding acquisition. All authors have read and approved the final manuscript.

## Acknowledgments

This work was supported by the National Natural Science Foundation of China (Grant Nos. 61972100 and 62172300).

## Supplementary material

**Figure S1.**
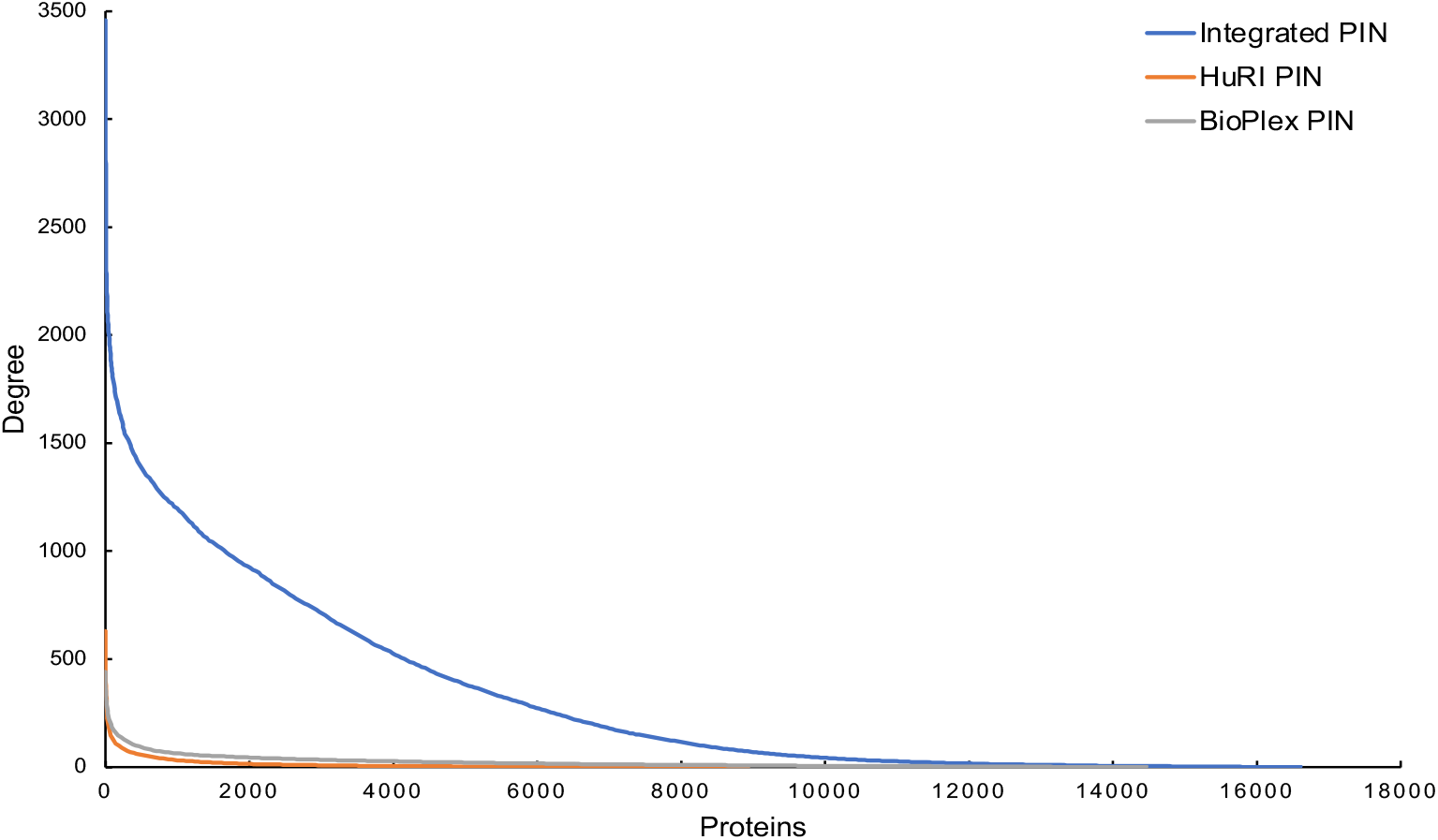
Degree distribution of proteins in different PINs. HuRI, BioPlex, and Integrated PIN are similar to scale-free networks.

**Figure S2.**
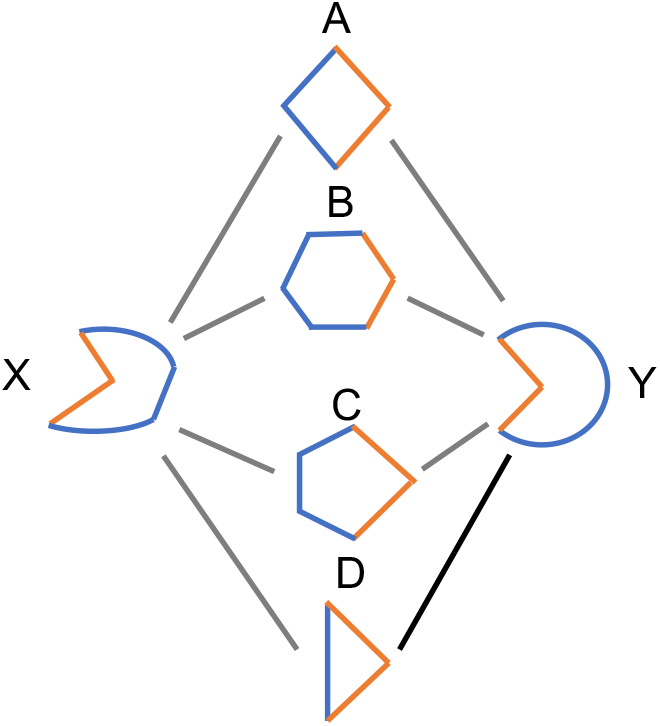
The L3 method diagram. PPIs often require complementary interfaces (e.g., protein C and protein Y). And protein D may interact with protein Y (black link), which can be predicted by using L3 method.

**Table S1.** The statistics of PINs.

| **PIN** | **No. of proteins** | **No. of PPIs** |
| --- | --- | --- |
| HuRI | 8985 | 63,132 |
| BioPlex | 14,484 | 167,932 |
| Integrated PIN | 16,632 | 2,658,160 |

*Note:* PPIs denotes protein-protein interactions.

**Table S2.**
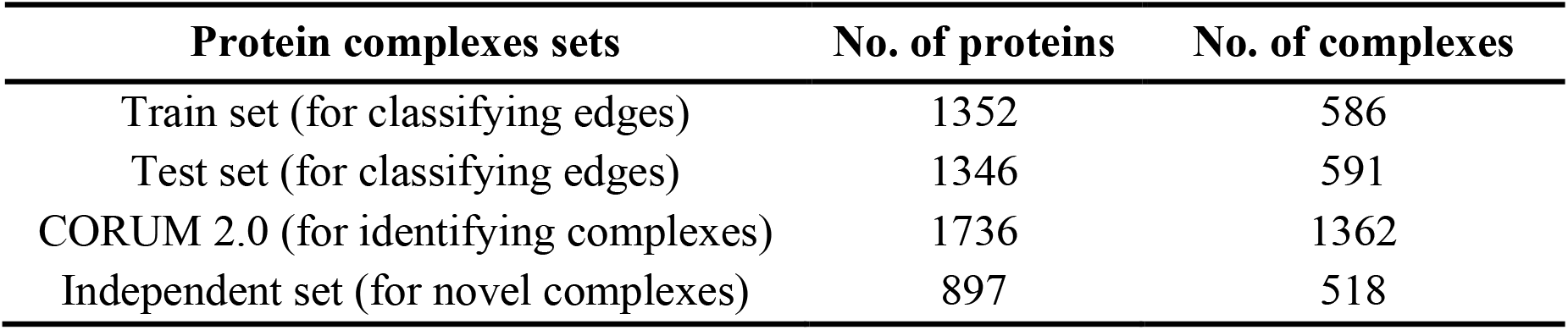
The statistics of protein complex sets.

| **Protein complexes sets** | **No. of proteins** | **No. of complexes** |
| --- | --- | --- |
| Train set (for classifying edges) | 1352 | 586 |
| Test set (for classifying edges) | 1346 | 591 |
| CORUM 2.0 (for identifying complexes) | 1736 | 1362 |
| Independent set (for novel complexes) | 897 | 518 |

**File S1 Algorithm for iteratively adding and removing nodes to generate a protein complex**

