## supplementary file 1 for "HPC-Atlas: Computationally Constructing A Comprehensive Atlas of Human Protein Complexes"

**Algorithm for iteratively adding and removing nodes to generate a protein complex**

**Algorithm 1 Iteratively generating a protein complex**

**Input:** The LE-PIN $G=(V,E,W)$,

${SG}_{v}$ is the subgraph that contains a single protein, from line 12 of article algorithm 1

**Output:** A protein complex, ${SG}_{v}$

1: **Initialize:** $k=1$ /* Record the number of iterations */
2: **Repeat**
3: **Step 1:** Add the neighbor nodes of the subgraph (${SG}_{v}$) to maximize the value of $F_{SG(v)}$ according to Equation (28)
4: **Initialize:** ${NSG}_{(v)}=\{\}$ /* The neighbor node set of the ${SG}_{v}$ */
5: Insert all neighbor nodes into ${NSG}_{(v)}$
6: **if** $\left| V_{{SG}_{v}} \right|=1$ **then**
7: $F_{SG(v)}=0$ and $Expectation\_edges=0$
8: **else**
9: Calculate the value of $F_{SG(v)}$ and $Expectation\_edges$ for ${SG}_{(v)}$ according to Equation (28) and (30)
10: **end if**
11: **while** $\left| {NSG}_{(v)} \right|\neq0$ **do**
12: **Initialize:** $node\_max=\emptyset$
13: **for** each protein $n_{s}$ in ${NSG}_{(v)}$ **do**
14: **if** $W(n_{s},V_{{SG}_{v}})>$ 0.5 **then** /* The edge weight of $n_{s}$ with the internal nodes of the ${SG}_{v}$ is greater than 0.5 */
15: Calculate the value of $F_{SG\left( v \right)+n_{s}}$for the subgraph (${SG}_{v}+n_{s}$) according to Equation (28)
16: Calculate the value of $Actually\_edges$ for the subgraph (${SG}_{v}+n_{s}$) according to Equation (31)
17: **if** $F_{SG\left( v \right)+n_{s}}> F_{SG(v)}$ and $Actually\_edges\geq Expectation\_edges$ **then**
18: $node\_max=n_{s}$; $F_{SG(v)}=F_{SG\left( v \right)+n_{s}}$;
19: $Expectation\_edges = Actually\_edges$
20: **end if**
21: **end if**
22: **end for**
23: **if** $node\_max\neq\emptyset$ **then**
24: Add $node\_max$ to ${SG}_{v}$; Remove $node\_max$ from ${NSG}_{(v)}$
25: **else**
26: break
27: **end if**
28: **end while**
29: **Step 2:** Remove any of inner nodes of the ${SG}_{v}$ to maximize the value of $F_{SG(v)}$
30: **Initialize:** ${BSG}_{v}=\{\}$ /* The boundary node set of the ${SG}_{v}$ */
31: **for** each protein $j$ in ${SG}_{v}$ **do**
32: Obtain the neighbor node set of $j$, denote as $Neighbor(j)$
33: $Interaction\_set = Neighbor(j)\cap V_{{SG}_{v}}$
34: **if**$Neighbor\left( j \right)-Interaction\_set\neq\emptyset$ **then** /* The difference set among $Neighbor(j)$ and $Interaction\_set$ */
35: Add $j$ into ${BSG}_{v}$
36: **end if**
37: **end for**
38: **while** $\left| {BSG}_{(v)} \right|\neq0$ **do**
39: **initialize:** $node\_min=\emptyset$
40: **for** each protein $n_{s}$ in ${BSG}_{(v)}$ **do**
41: Calculate the value of $F_{SG\left( v \right)-n_{s}}$ for the subgraph (${SG}_{v}-n_{s}$) according to Equation (28)
42: Calculate the value of $Actually\_edges$ for the subgraph (${SG}_{v}-n_{s}$) according to Equation (31)
43: **if** $F_{SG\left( v \right)-n_{s}}> F_{SG(v)}$ and $Actually\_edges\geq Expectation\_edges$ **then**
44: $node\_min=n_{s}$; $F_{SG(v)}=$ $F_{SG\left( v \right)-n_{s}}$;
45: $Expectation\_edges = Actually\_edges$
46: **end if**
47: **end for**
48: **if** $node\_min\neq\emptyset$ **then**
49: Remove $node\_min$ from ${SG}_{v}$; Remove $node\_min$ from ${BSG}_{(v)}$
50: **else**
51: break
52: **end if**
53: **end while**
54: $k=k+1$
55: **until** ${SG}_{v}$ is not changing
56: **return** Output the predicted protein complex, ${SG}_{v}$
